# Polygenic Risk Score for Smoking is associated with Externalizing Psychopathology and Disinhibited Personality Traits but not Internalizing Psychopathology in Adolescence

**DOI:** 10.1101/2020.07.29.227405

**Authors:** Brian M. Hicks, D. Angus Clark, Joseph D. Deak, Mengzhen Liu, C. Emily Durbin, Jonathan D. Schaefer, Sylia Wilson, William G. Iacono, Matt McGue, Scott I. Vrieze

**Affiliations:** Department of Psychiatry, University of Michigan; Department of Psychiatry, Yale University; Department of Psychology, University of Minnesota; Department of Psychology, Michigan State University; Institute of Child Development, University of Minnesota

## Abstract

**Importance:** Large consortia of genome wide association studies have yielded more accurate polygenic risk scores (PRS) that aggregate the small effects of many genetic variants to characterize the genetic architecture of disorders and provide a personalized measure of genetic risk.

**Objective:** We examined whether a PRS for smoking measured genetic risk for general behavioral disinhibition by estimating its associations with externalizing and internalizing psychopathology and related personality traits. We examined these associations at multiple time points in adolescence using more refined phenotypes defined by stable characteristics across time and at young ages, which reduced potential confounds associated with cumulative exposure to substances and reverse causality.

**Methods:** Random intercept panel models were fit to symptoms of conduct disorder, oppositional defiant disorder, major depressive disorder (MDD), and teacher ratings of externalizing and internalizing problems and personality traits at ages 11, 14, and 17 years-old in the Minnesota Twin Family Study (*N* = 3225).

**Results:** The smoking PRS had strong associations with the random intercept factors for all the externalizing measures (mean standardized *β* = .27), agreeableness (*β=*−.22, 95% CI: −.28, −.16), and conscientiousness (*β=*−.19, 95% CI: −.24, −.13), but was not significantly associated with the internalizing measures (mean *β* = .06) or extraversion (*β=*.01, 95% CI: −.05, .07). After controlling for smoking at age 17, the associations with the externalizing measures (mean *β* = .13) and personality traits related to behavioral control (mean *β* = −.10) remained statistically significant.

**Conclusions and Relevance:** The smoking PRS measures genetic influences that contribute to a spectrum of phenotypes related to behavioral disinhibition including externalizing psychopathology and normal-range personality traits related to behavioral control, but not internalizing psychopathology. Continuing to identify the correlates and delineate the mechanisms of the genetic influences associated with disinhibition could have substantial impact in mitigating a variety of public health problems (e.g., mental health, academic achievement, criminality).

**Key Points:** *Question:* Does a polygenic risk scores (PRS) for smoking measure genetic risk for behavioral disinhibition in general?

*Findings:* The smoking PRS was associated with externalizing psychopathology and personality traits related to behavioral control, but not internalizing psychopathology and extraversion during adolescence, even after controlling for smoking status.

*Meaning:* The smoking PRS measures genetic influences on behavioral disinhibition in general which is associated with a variety of important outcomes including mental health, academic success, and criminality.

## Introduction

Substance use disorders (SUDs) are highly comorbid with each other, antisocial behavior, and normal-range personality traits associated with disinhibition (e.g., impulsivity, antagonism)(1-3). Externalizing problems (e.g., rule breaking, aggression), in particular, are early emerging behaviors that are robust predictors of later SUDs (4), as are parental externalizing disorders (5). Multivariate behavior genetic studies indicate that a broad, highly heritable liability to behavioral disinhibition best accounts for the associations among SUDs, antisocial behavior, and disinhibited personality traits, as well as their familial transmission (2, 5-7). Although internalizing problems (depression, anxiety) are also associated with SUDs (1, 8) relative to externalizing problems, these associations are smaller, less robust in terms of being early predictors of later SUDs, and there is less evidence of shared genetic influences (8-10)

Large consortia of genome wide association studies (GWAS) have detected hundreds of genome-wide significant (*p* = 5.0×10^−8^) associations between specific genetic variants and psychopathology. However, effect sizes for specific variants are typically very small; therefore, the cumulative effect of either all or a subset (e.g., those of genome-wide significance) of measured genetic variants on a phenotype is often estimated using polygenic risk scores (PRSs). PRSs are calculated by first weighting genetic variants based on the magnitude of their association with a phenotype of interest in a discovery sample, and then summing these weighted allele counts in a target sample to examine the association between the “risk score” and outcomes in the target sample.

One useful application of PRSs is the ability to draw the boundaries of genetic influences on psychiatric disorders. For example, there is significant overlap among PRS for schizophrenia, bipolar, autism spectrum disorder, attention deficit hyperactivity disorder, major depression (11, 12), and among alcohol, nicotine, and cannabis use (13-15), as well as some evidence of genetic specificity (16). Relatedly, childhood symptoms are associated with PRS for adult disorders, indicating that these genetic influences contribute to the expression of psychiatric problems early in life. For example, a meta-analysis of seven longitudinal studies found that the combined symptoms of ADHD, internalizing, and social problems were associated with PRSs derived in adult samples for major depression, neuroticism, low subjective well-being, insomnia, low educational attainment, and body mass index (17).

Our goal was to clarify the nomological network of a PRS for a binary phenotype of having ever smoked regularly that was derived from the largest GWAS of smoking-related phenotypes to date (*N*=1,232,091), which yielded 378 genome-wide significant hits (13). In replication samples, this PRS accounted for 4% of the phenotypic variance in smoking and was associated with use of a variety of other substances (e.g., alcohol, cannabis, cocaine, amphetamines, ecstasy, hallucinogens) (13, 18). Using the same sample as in the current report, we also found that this smoking PRS predicted trajectories of nicotine and alcohol use from ages 14 to 34. Notably, the smoking PRS was a stronger predictor of both nicotine *and* alcohol use trajectories than a PRS for number of alcoholic drinks per week, even after accounting for the genetic correlation between the two PRSs and the phenotypic overlap between nicotine and alcohol use (19). Finally, this smoking PRS was associated with the externalizing dimension of the Child Behavior Checklist in a large sample of pre-adolescents, even after adjusting for the general factor of psychopathology (20). These findings suggest the smoking PRS may measure genetic risk for behavioral disinhibition in general rather than smoking specifically.

To test this hypothesis, we examined the associations between multiple measures of externalizing and internalizing psychopathology and related personality traits. Importantly, we took a developmental approach, and focused on behavioral phenotypes that were assessed on multiple occasions at young ages (11, 14, and 17-years old). This allowed us to define each phenotype by characteristics that were stable across time (i.e., without time-specific influences and unsystematic measurement error), and reduced the potential for reverse causality due to the negative consequences of heavy substance use (e.g., smoking PRS → smoking → externalizing). We predicted that the smoking PRS would be associated with externalizing problems, but have smaller or null associations with internalizing problems. We also predicted that the smoking PRS would be associated with normative manifestations of behavioral disinhibition, specifically, personality traits associated with behavioral control (i.e., agreeableness and conscientiousness), but have smaller or null associations with traits associated with emotionality (i.e., neuroticism and extraversion).

## Methods

### Participants

Participants were members of the Minnesota Twin Family Study (MTFS), a longitudinal study of 3762 (52% female) twins (1881 pairs) investigating the development of substance use disorders and related conditions (21-23). All twin pairs were the same sex and lived with at least one biological parent within driving distance to the University of Minnesota laboratories at the time of recruitment. Exclusion criteria included any cognitive or physical disability that would interfere with study participation. Twins were recruited the year they turned either 11-years old (*n* = 2510; the younger cohort) or 17-years old (*n* = 1252; the older cohort). Twins in the younger cohort were born from 1977 to 1984 and 1988 to 1994, while twins in the older cohort were born between 1972 and 1979. Families were representative of the area they were drawn from in terms of socioeconomic status, history of mental health treatment, and urban vs rural residence (21). Consistent with the demographics of Minnesota for the target birth years, 96% of participants reported European American ancestry.

The younger cohort was assessed at ages 11 (M_age_ = 11.78 years; SD = 0.43 years) and 14 (M_age_ = 14.90 years; SD = 0.31 years), and all twins were assessed at age 17 (M_age_ = 17.85 years; SD = 0.64 years). Table 1 provides the number of participants for each assessment and descriptive statistics for the study measures. Retention rates were 91.4% and 86.3% at ages 14 and 17, respectively, for twins in the younger cohort. The total sample included 1205 monozygotic (51.5% female) and 676 dizygotic (52.8% female) twin pairs.

**Table 1.** Descriptive Information for Outcome Variables Across Time

|  | Age 11 |  |  | Age 14 |  |  |  | Age 17 |  |  |  |  |
| --- | --- | --- | --- | --- | --- | --- | --- | --- | --- | --- | --- | --- |
|  | M | SD | N | M | SD | N | *r*11 | M | SD | N | *r*11 | *r*14 |
| **Externalizing Psychopathology** |  |  |  |  |  |  |  |  |  |  |  |  |
| Externalizing Factor | 0 | .85 | 2510 | 0 | .88 | 2345 | .60 | 0 | .81 | 3512 | .48 | .65 |
| Conduct Disorder | 0 | 1 | 2510 | 0 | 1 | 2291 | .45 | 0 | 1 | 3414 | .30 | .51 |
| Oppositional Defiant Disorder | 2.10 | 1.87 | 2510 | 2.44 | 2.03 | 2291 | .49 | 2.37 | 1.98 | 3468 | .40 | .57 |
| Teacher Rated Externalizing Problems | 1.40 | .46 | 2292 | 1.35 | .43 | 1931 | .62 | 1.44 | .46 | 2664 | .55 | .64 |
| **Internalizing Psychopathology** |  |  |  |  |  |  |  |  |  |  |  |  |
| Major Depressive Disorder | .34 | 1.22 | 2506 | .55 | 1.62 | 2289 | .15 | 1.03 | 2.26 | 3503 | .17 | .32 |
| Teacher Rated Internalizing Problems | 1.59 | .45 | 1820 | 1.53 | .45 | 1901 | .29 | 1.55 | .45 | 2375 | .33 | .36 |
| **Teacher Rated Personality Traits** |  |  |  |  |  |  |  |  |  |  |  |  |
| Extraversion | 3.21 | .70 | 2292 | 3.20 | .70 | 1932 | .52 | 3.35 | .66 | 2672 | .47 | .54 |
| Neuroticism | 1.74 | .71 | 2291 | 1.70 | .67 | 1928 | .37 | 1.76 | .68 | 2662 | .30 | .37 |
| Conscientiousness | 3.39 | .80 | 2291 | 3.53 | .77 | 1933 | .61 | 3.64 | .72 | 2672 | .56 | .59 |
| Agreeableness | 3.94 | .77 | 2291 | 3.96 | .73 | 1929 | .52 | 3.98 | .70 | 2670 | .46 | .53 |

M = mean; SD = standard deviation; N = number of ratings; *r*11 = correlation with corresponding variable at age 11; *r*14 = correlation with corresponding variable at age 14. Because conduct disorder symptoms must be present prior to age 15, the adult criteria for antisocial personality disorder was used to assess conduct disorder at age 17. Consequently, values for conduct disorder symptom counts were standardized within each time point. Externalizing factor scores were derived from time-point specific factor models with conduct disorder, oppositional defiant disorder, and teacher rated externalizing problems as indicators. Factor means and variances were fixed to 0 and 1, respectively. Due to the within-time standardization of conduct disorder and specification of the externalizing factor there is no mean change over time apparent in these variables; as the primary analyses do not draw on the mean structure though this does not affect the major results.

### Conduct Disorder (CD), Adult Antisocial Behavior (AAB), Oppositional Defiant Disorder (ODD), and Major Depressive Disorder (MDD)

Structured clinical interviews were used to obtain symptom counts for CD, ODD, and MDD using *DSM-III-R* criteria, which was the diagnostic system used when the study began. Because CD symptoms must be present before age 15, the adult criteria for antisocial personality disorder was used to assess antisocial behavior at age 17. Twins and their mothers reported on symptoms of CD, ODD, and MDD. A symptom was considered present if reported by either informant. The reporting period for symptoms was lifetime for the age 11 assessment and the age 17 assessment for the older cohort, and the last 3-years at the age 14 and 17 assessments for the younger cohort. In combination, these symptom assessments take advantage of the best interview data at each age, optimizing the capture of diagnostic information available through the course of adolescence. Twins also reported on their quantity and frequency of nicotine use as well as symptoms of nicotine dependence during a structured interview, which were used to calculate a composite measure of smoking at age 17 (19).

### Teacher Ratings of Externalizing and Internalizing Problems

A teacher rating form covering various aspects of personality, behavior, and adjustment was completed by up to 3 teachers nominated by the twin participants (4). The mean rating across teachers was used whenever more than one teacher rating was available (∼75% of participants with teacher rating data had at least two teacher informants). It was Minnesota state policy to place members of twin pairs in separate classrooms whenever possible, which minimizes any bias due to twin contrast or comparison.

Teachers were instructed to rate how characteristic specific problem behaviors were of each student (1 = *not at all*, 2 = *just a little*, 3 = *pretty muc*h, 4 = *very much*). A teacher rating of externalizing (TR-EXT) problems was calculated using 28-items (mean α = .97) related to inattention, oppositionality and defiance, impulsivity, and aggression, while a teacher rating of internalizing (TR-INT) problems was calculated using 8-items (mean α = .83) related to feelings of distress, unhappiness, fear, anxiety, and stress reactivity.

### Teacher Ratings of Personality

The teacher rating form also included 28 items that consisted of short descriptors designed to be markers of personality trait constructs (24). For each item, teachers were instructed to rank each twin in reference to their classmates as falling in either the *lowest 5%, lower 30%, middle 30%, higher 30%*, or *highest 5%* of students in class. Exploratory and confirmatory factor analyses were used to construct four scales that corresponded to trait constructs in Big Five trait models of personality: extraversion (6-items: mean α = .86), neuroticism (4-items: mean α = .76), conscientiousness (5-items: mean α = .91), agreeableness/prosociality (5-items: mean α = .89).

### PRS Methods

PRSs were generated from the GSCAN discovery sample using GWAS summary statistics for having ever smoked regularly, following removal of the MTFS sample to guard against overlap with the target sample (13). PRSs were created for participants of European ancestry in the MTFS target sample following imputation to the most recent Haplotype Reference Consortium reference panel (25), and restricted to variants with a minor allele frequency ≥ 0.001. The resulting filtered variants (i.e., 2.8 million variants) were then submitted to LDpred (26) to generate beta weights in the MCTFR target sample, including variants of all significance levels (i.e., *p*-value threshold ≤ 1) in order to capture all degrees of genetic influences across the genome. Individual PRSs were then calculated in PLINK 1.9 (27) for all individuals meeting inclusion criteria for the present study (*N* = 3225).

### Data Analytic Strategy

Associations between the regular smoking PRS and the longitudinal measures of psychopathology and personality were examined using multiple regression and random intercept panel models (RI-PM) (see Figure 1). Externalizing factor scores were also calculated using the covariance among the available measures at each time point (CD, ODD, TR-EXT). In the multiple regression models, a psychopathology or personality variable at a single time point was entered as the outcome and regressed on the smoking PRS and the covariates of participant sex and the first five genetic principal components to adjust for ancestral stratification.

**Figure 1.**
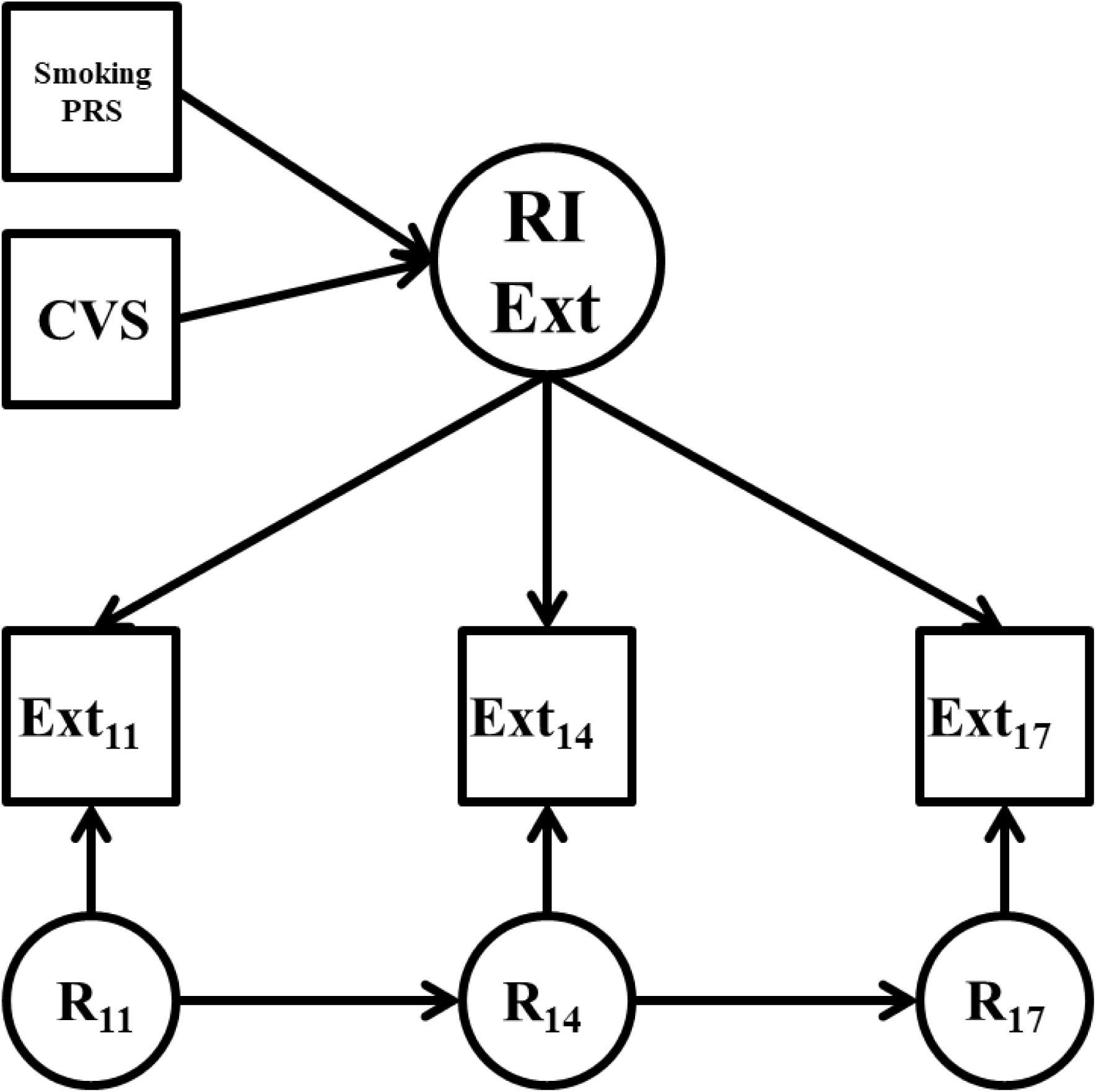
Conditional Random Intercept Model for Externalizing Problems. RI = random intercept factor; EXT = externalizing at ages 11, 14, and 17; R = residual factors at age 11, 14, and 17; CVS = set of covariates including the first five genetic principal components, sex, and occasionally the nicotine use composite at age 17. Variances/residual variances and mean structure omitted from figure for clarity of presentation.

For the RI-PMs, measures of psychopathology or personality (Ext_11_, Ext_14_, Ext_17_ in Figure 1) at each time point were specified to load on a time-invariant random intercept factor (RI Ext in Figure 1). Factor loadings were fixed to 1, and indicator intercepts were allowed to vary. The mean of the random intercept was fixed to 0 and the variance was freely estimated. The random intercept captures the common trait variance across time (28). For example, a positive random intercept score indicates an individual consistently ranked higher than the mean of their peers across time. Occasion-specific residual factors (R_11_ through R_17_ in Figure 1) were then specified with factor loadings fixed to 1, means fixed to 0, and the variances were freely estimated. Autoregressive paths were also added from one residual factor to the subsequent residual factor (αx in Figure 1), to account for correlations between time-point specific deviations from the overall trend line. The random intercept factors were regressed on the smoking PRS and the covariates of participant sex and the first five genetic principal components (29), as well as smoking at age 17 to test whether the smoking PRS remained a significant predictor after adjusting for the putatively expressed phenotype.

All models were fit in Mplus version 8.4 (30) using full information maximum likelihood estimation. Confidence intervals were derived via clustered (by family) nonparametric percentile bootstrap (1000 draws), which provides reliable assessments of parameter estimate precision under a variety of complex data conditions (31). A parameter estimate was considered statistically significant if the bootstrapped 95% confidence interval did not include 0. The Mplus Automation Package (32) in R (33) was also used to facilitate the analyses.

## Results

Descriptive statistics for the study variables are reported in Table 1. The externalizing (mean autocorrelation = .52) and personality (mean autocorrelation = .49) measures had moderate to strong stability over time, while the internalizing measures had only small to moderate stability over time (mean autocorrelation = .27). The unconditional RI-PMs were fully saturated and thus were a perfect to the data. The variance component of each random intercept was statistically significant (see Table 2 for parameter estimates). Results for the multiple regression and RI-PMs for the youth externalizing, internalizing, and personality measures are presented in Table 3.

**Table 2.** Unconditional Random Intercept Model Results

Unconditional Random Intercept Model Results
| | $\sigma_{ri}$ | $\sigma_{r11}$ | $\psi_{r14}$ | $\psi_{r17}$ | $r11 \rightarrow r14$ | $r14 \rightarrow r17$ |
| --- | --- | --- | --- | --- | --- | --- |
| **Externalizing Psychopathology** |  |  |  |  |  |  |
| Externalizing Factor | .25* | .46* | .43* | .32* | .41* | .41* |
| Conduct Disorder | .21* | .79* | .70* | .69* | .30* | .38* |
| Oppositional Defiant Disorder | 1.23* | 2.28* | 2.76* | 2.30* | .29* | .37* |
| Teacher Rated Externalizing Problems | .12* | .10* | .07* | .10* | .05 | .15 |
| **Internalizing Psychopathology** |  |  |  |  |  |  |
| Major Depressive Disorder | .53* | .96* | 2.05* | 4.41* | -.25 | .30* |
| Teacher Rated Internalizing Problems | .07* | .13* | .14* | .14* | -.08 | .05 |
| **Teacher Rated Personality Traits** |  |  |  |  |  |  |
| Extraversion | .21* | .28* | .26* | .22* | .15* | .12 |
| Neuroticism | .14* | .36* | .30* | .33* | .11* | .10 |
| Conscientiousness | .32* | .32* | .27* | .21* | .18* | .04 |
| Agreeableness | .25* | .33* | .28* | .26* | .11 | .09 |

$\sigma_{ri}$ = random intercept variance; $\sigma_{r11}$ = age 11 residual factor variance; $\psi_{r14}$ = age 14 residual factor residual variance; $\psi_{r17}$ = age 17 residual factor residual variance; $r11 \rightarrow r14$ = autoregressive path from age 11 residual factor to age 14 residual factor; $r14 \rightarrow r17$ = autoregressive path from age 14 residual factor to age 17 residual factor; \* = 95% confidence interval for parameter estimate did not include 0. Unstandardized parameter estimates presented. All models were fully saturated (i.e., degrees of freedom = 0) and thus fit perfectly. Confidence intervals derived via clustered, non-parametric percentile bootstrapping (with 1,000 random draws).

**Table 3.** Standardized Regression Coefficients From Smoking PRS to Outcome Variables

|  | Age 11 | Age 14 | Age 17 | Random Intercept Factor<br>adj for covs | adj for covs &<br>smoking |
| --- | --- | --- | --- | --- | --- |
| **Externalizing Psychopathology** |  |  |  |  |  |
| Externalizing Factor | **.22**<br>[.16,.27] | **.23**<br>[.17,.28] | **.24**<br>[.18,.28] | **.30**<br>[.24,.37] | **.14**<br>[.09,.19] |
| Conduct Disorder | **.16**<br>[.11,.21] | **.16**<br>[.11,.21] | **.18**<br>[.14,.22] | **.36**<br>[.28,.56] | **.15**<br>[.10,.20] |
| Oppositional Defiant Disorder | **.14**<br>[.08,.19] | **.18**<br>[.13,.23] | **.16**<br>[.11,.20] | **.26**<br>[.20,.34] | **.15**<br>[.10,.22] |
| Teacher Rated Externalizing Problems | **.14**<br>[.08,.19] | **.17**<br>[.12,.23] | **.09**<br>[.05,.14] | **.15**<br>[.10,.20] | **.08**<br>[.03,.13] |
| **Internalizing Psychopathology** |  |  |  |  |  |
| Major Depressive Disorder | <.01<br>[-.06,.04] | .04<br>[-.01,.09] | **.08**<br>[.04,.12] | .07<br>[-.01,.15] | .01<br>[-.06,.09] |
| Teacher Rated Internalizing Problems | <.01<br>[-.07,.06] | .03<br>[-.03,.09] | .02<br>[-.22,.07] | .04<br>[-.03,.10] | -.01<br>[-.07,.06] |
| **Teacher Rated Personality Traits** |  |  |  |  |  |
| Extraversion | <.01<br>[-.06,.06] | .02<br>[-.04,.07] | .02<br>[-.03,.07] | .01<br>[-.05,.07] | .03<br>[-.03,.09] |
| Neuroticism | **.08**<br>[.02,.13] | **.07**<br>[.01,.12] | **.07**<br>[.02,.12] | **.14**<br>[.07,.21] | .04<br>[-.02,.10] |
| Conscientiousness | **-.13**<br>[-.18,-.07] | **-.17**<br>[-.22,-.12] | **-.13**<br>[-.18,-.08] | **-.19**<br>[-.24,-.13] | **-.10**<br>[-.15,-.06] |
| Agreeableness | **-.12**<br>[-.12,-.06] | **-.17**<br>[-.23,-.13] | **-.16**<br>[-.20,-.11] | **-.22**<br>[-.28,-.16] | **-.10**<br>[-.15,-.05] |

adj for covs = regression paths from smoking PRS to random intercept adjusted for standard covariates; adj for covs & smoking = regression paths from smoking PRS to random intercept after adjusting for standard covariates and smoking at age 17; Age 11, 14, and 17 = regression paths from smoking PRS to outcome variables at ages 11, 14, and 17, respectively. All models include the standard covariates of participant sex and five genetic principal components to adjust for ancestry stratification. 95% confidence intervals presented under parameter estimates. Parameter estimates for which confidence intervals do not include 0 are presented in bold. Confidence intervals derived via clustered, non-parametric percentile bootstrapping (with 1,000 random draws).

### Adolescent Externalizing Problems

For the externalizing measures, all the regression coefficients were statistically significant and small to medium in size (mean standardized *β* = .17). In the RI-PMs, the smoking PRS had small to medium associations with the random intercept factors for each externalizing measure (mean *β* = .27). The associations with the random intercept factors remained significant after adjusting for smoking at age 17, though the effect sizes were reduced by about 50% (mean *β* = .13).

### Adolescent Internalizing Problems

In contrast to the externalizing phenotypes, associations between the internalizing measures and the smoking PRS were null to small in size (mean *β* = .03), and only the association with MDD at age 17 was statistically significant (*β* = .08, 95% CI: .04, .12). The smoking PRS had small and non-significant associations with the random intercept factors for MDD (*β* = .07, 95% CI: −.01, .15) and TR-INT problems (*β* = .04, 95% CI: −.03, .10), and adjusting for smoking at age 17 reduced the magnitude of these associations to zero.

### Adolescent Personality

The regression coefficients for neuroticism, conscientiousness, and agreeableness were all statistically significant and small to medium in size (mean |*β*| = .12). The smoking PRS had small to medium associations with the random intercept factors for agreeableness (*β* = −.22, 95% CI: −.28, −.16), conscientiousness (*β* = −.19, 95% CI: −.24, −.13), and neuroticism (*β* = .14, 95% CI: .07, .21). Adjusting for smoking at age 17 reduced the associations with agreeableness and conscientiousness by about 50%, though the adjusted associations remained statistically significant. In contrast, the association with neuroticism declined by about 70% and was no longer significant. The associations between the smoking PRS and the measures of extraversion and its random intercept factor were near zero and not statistically significant (mean |*β*| = .01).

## Discussion

The results provided strong evidence that the smoking PRS measures genetic risk for behavioral disinhibition, as evidenced by the consistent and relatively strong associations with CD, ODD, and TR-EXT at multiple time points and across different informants from ages 11 to 17. These associations were stronger for the variance that was stable across time (i.e., the random intercept factor), suggesting that these genetic variants contribute to consistent individual differences in psychological processes that contribute to externalizing problems over time. The genetic influences measured by the smoking PRS extended beyond symptoms of pathology to normal range personality traits associated with behavioral control (conscientiousness and agreeableness), indicating this aggregate genetic risk also contributes to broad temperament dispositions.

Furthermore, these associations remained significant—though were reduced in size— after controlling for the putatively expressed phenotype of smoking at age 17, which helps rule out the possibility of reverse causation (i.e., smoking PRS → smoking → externalizing). Conversely, the smoking PRS was *not* associated with internalizing problems or extraversion, and its association with neuroticism dropped to non-significance after adjusting for smoking at age 17. Taken together, these results exhibit a theoretically coherent pattern of associations indicating the smoking PRS indexes genetic risk for behavioral disinhibition, but not psychopathology in general. This is consistent with multivariate twin studies that have posited a common genetic etiology that accounts for the co-occurrence among SUDs, antisocial behavior, and disinhibited personality traits (2, 5-7)

Behavioral disinhibition is associated with a variety of outcomes (34), and it will be important to continue to test if the correlates of the smoking PRS maps onto the broader nomological network of behavioral disinhibition. For example, academic engagement and success and criminality are especially important domains to examine further. The smoking PRS also has the potential to help delineate processes of gene-environment correlation and interaction for important outcomes. If the empirical correlates of the smoking PRS closely align with the nomological network of behavioral disinhibition, it may also help to delineate the mechanisms that underlie the psychological processes of disinhibition. A limitation of the current PRS method, however, is that it is a crude estimate of overall genetic influence that does not identify specific variants that point to specific biological processes. A key next step will be to incorporate functional genomic information (35, 36) to begin the process of uncovering the neurobehavioral mechanisms that account for the link between genetic background and clinical phenotypes. Another goal is that PRS approaches will eventually have practical value by identifying persons at greatest risk for experiencing negative outcomes, and will inform prevention and intervention efforts (37). PRSs that have been validated with longitudinal data may be especially helpful in identifying high-risk individuals at young ages to circumvent negative outcomes, and in crafting PRS-informed interventions that forestall the development of severe psychopathology that may persist through adulthood (38).

The study has some limitations including a sample was restricted to people of European ancestry. Consequently, it is unclear if the predictive utility of the smoking PRS will generalize to people of other ancestral groups due to varying allele frequencies (39, 40). Notably, this limitation has the potential to proliferate health disparities with precision medicine efforts if these findings are only applicable to individuals of European ancestry, further prioritizing the importance of extending these efforts to diverse ancestry groups (41). Additionally, genetic influences always act within an environmental context, and several putatively environmental variables such as peers and parenting have strong associations with behavioral disinhibition, the effects of which may vary across development (42, 43). Delineating the gene-environment interplay between disinhibition and known environmental risk factors may be key reaping practical utility from the smoking PRS.

Despite these limitations, the current results extend prior work by validating the predictive utility of the smoking PRS as a genetic measure of general behavioral disinhibition. This validation serves as a key step in demonstrating that polygenic prediction of disinhibition-related outcomes across important developmental periods has the potential to have meaningful implications for the future of personalized prevention and intervention efforts for a variety of consequential outcomes (e.g., education, work, mental health, criminality).

## Acknowledgement

This work was supported by United States Public Health Service grants R01 AA024433 (Hicks), R01 AA09367 (McGue), R21 AA026632 (Wilson), and T32 AA007477 (Blow) from the National Institute on Alcohol Abuse and Alcoholism; R01 DA034606 (Hicks), R01 DA042755 (Hopfer), R01 DA013240 (Iacono), R37 DA005147 (Iacono), R01 DA044283 (Vrieze), R01 DA037904 (Vrieze), U01 DA046413 (Vrieze), and K01 DA037280 (Wislon) from the National Institute on Drug Abuse; T32 MH015755 (Cicchetti) from the National Institute of Mental Health.

## References

1. Kessler RC, Chiu WT, Demler O, Walters EE. Prevalence, severity, and comorbidity of 12-month DSM-IV disorders in the National Comorbidity Survey Replication. Arch Gen Psychiatry. 2005;62(6):617–27.

2. Krueger RF, Hicks BM, Patrick CJ, Carlson SR, Iacono WG, McGue M. Etiologic connections among substance dependence, antisocial behavior, and personality: Modeling the externalizing spectrum. Journal of Abnormal Psychology. 2002;111(3):411–24.

3. Agrawal A, Dick DM, Bucholz KK, Madden PAF, Cooper ML, Sher KJ, et al. Drinking expectancies and motives: a genetic study of young adult women. Addiction. 2008;103(2):194–204.

4. Hicks BM, Iacono WG, McGue M. Identifying childhood characteristics that underlie premorbid risk for substance use disorders: Socialization and boldness. Dev Psychopathol. 2014;26(1):141–57.

5. Hicks BM, Foster KT, Iacono WG, McGue M. Genetic and Environmental Influences on the Familial Transmission of Externalizing Disorders in Adoptive and Twin Offspring. Jama Psychiatry. 2013;70(10):1076–83.

6. Kendler KS, Prescott CA, Myers J, Neale MC. The structure of genetic and environmental risk factors for common psychiatric and substance use disorders in men and women. Archives of General Psychiatry. 2003;60(9):929–37.

7. Young SE, Stallings MC, Corley RP, Krauter KS, Hewitt JK. Genetic and environmental influences on behavioral disinhibition. American Journal of Medical Genetics. 2000;96(5):684–95.

8. Tully EC, Iacono, W. G. An integrative common liabilities model for the comorbidity of substance use disorders with externalizing and internalizing disorders. In: Sher KJ, editor. The Oxford handbook of substance use and substance use disorders. Oxford: Oxford University Press; 2016. p. 187–212.

9. Foster KT, Hicks BM, Zucker RA. Positive and Negative Effects of Internalizing on Alcohol Use Problems From Childhood to Young Adulthood: The Mediating and Suppressing Role of Externalizing. Journal of Abnormal Psychology. 2018;127(4):394–403.

10. Saraceno L, Munafo M, Heron J, Craddock N, van den Bree MBM. Genetic and non-genetic influences on the development of co-occurring alcohol problem use and internalizing symptomatology in adolescence: a review. Addiction. 2009;104(7):1100–21.

11. Smoller JW, Craddock N, Kendler K, Lee PH, Neale BM, Nurnberger JI, et al. Identification of risk loci with shared effects on five major psychiatric disorders: a genome-wide analysis. Lancet. 2013;381(9875):1371–9.

12. Lee SH, Ripke S, Neale BM, Faraone SV, Purcell SM, Perlis RH, et al. Genetic relationship between five psychiatric disorders estimated from genome-wide SNPs. Nature Genet. 2013;45(9):984-+.

13. Liu MZ, Jiang Y, Wedow R, Li Y, Brazel DM, Chen F, et al. Association studies of up to 1.2 million individuals yield new insights into the genetic etiology of tobacco and alcohol use. Nature Genet. 2019;51(2):237-+.

14. Quach BC, Bray, M. J., Gaddis, N. C., Liu, M., Palviainen, T., Minica, C. C., Zellers, S., Sherva, R., Aliev, F., & Nothnagel, M. Expanding the genetic architecture of nicotine dependence and its shared genetics with multiple traits: Findings from the Nicotine Dependence GenOmics (iNDiGO) Consortium. BioRxiv. 2020.

15. Zhou H, Sealock, J. M., Sanchez-Roige, S., Clarke, T.-K., Levey, D., Cheng, Z., Li, B., Polimanti, R., Kember, R. L., & Smith, R. V. Meta-analysis of problematic alcohol use in 435,563 individuals identifies 29 risk variants and yields insights into biology, pleiotropy and causality. BioRxiv, 738088. 2019.

16. Byrne EM, Zhu ZH, Qi T, Skene NG, Bryois J, Pardinas AF, et al. Conditional GWAS analysis to identify disorder-specific SNPs for psychiatric disorders. Mol Psychiatr. 12.

17. Akingbuwa WA, Hammerschlag, A. R., Jami, E. S., et al., Bipolar Disorder and Major Depressive Disorder Working Groups of the Psychiatric Genomics Consortium. Genetic associations between childhood psychopathology and adult depression and associated traits in 42,998 individuals: A meta-analysis. JAMA Psychiatry. 2020;77:715–28.

18. Chang LH, Couvy-Duchesne B, Liu MZ, Medland SE, Verhulst B, Benotsch EG, et al. Association between polygenic risk for tobacco or alcohol consumption and liability to licit and illicit substance use in young Australian adults. Drug Alcohol Depend. 2019;197:271–9.

19. Deak JD, Clark, D. A., Liu, M., Durbin, C. E., Iacono, W. G., McGue, M., Vrieze, S. I., Hicks, B. M. Polygenic risk scores predict the development of alcohol and nicotine use problems from adolescence through young adulthood. https://doiorg/1031234/osfio/axkqb. 2020, July 2.

20. Waszczuk MA, Miao, J., Docherty, A. R., Shabaliln, A. A., Jonas, K. G., Michelini, G., & Kotov, R. General vs. specific vulnerabilities: Polygenic risk scores and higher-order psychopathology dimensions in the Adolescent Brain Cognitive Development (ABCD) Study https://doiorg/1031234/osfio/km6v3. 2020, May 22.

21. Iacono WG, Carlson SR, Taylor J, Elkins IJ, McGue M. Behavioral disinhibition and the development of substance-case disorders: Findings from the Minnesota Twin Family Study. Development and Psychopathology. 1999;11(4):869–900.

22. Keyes MA, Malone SM, Elkins IJ, Legrand LN, McGue M, Iacono WG. The Enrichment Study of the Minnesota Twin Family Study: Increasing the yield of twin families at high risk for externalizing psychopathology. Twin Research and Human Genetics. 2009;12(5):489–501.

23. Wilson S, Haroian K, Iacono WG, Krueger RF, Lee JMJ, Luciana M, et al. Minnesota Center for Twin and Family Research. Twin Res Hum Genet. 2019;22(6):746–52.

24. Clark DA, Durbin, C. E., Iacono, W. G., McGue, M., & Hicks, B. M. Personality and sexual development in adolescence: Selection, corresponsive effects, and genetic and environmental influences. 2020.

25. Das S, Forer L, Schonherr S, Sidore C, Locke AE, Kwong A, et al. Next-generation genotype imputation service and methods. Nature Genet. 2016;48(10):1284–7.

26. Vilhjalmsson BJ, Yang J, Finucane HK, Gusev A, Lindstrom S, Ripke S, et al. Modeling Linkage Disequilibrium Increases Accuracy of Polygenic Risk Scores. Am J Hum Genet. 2015;97(4):576–92.

27. Chang CC, Chow CC, Tellier L, Vattikuti S, Purcell SM, Lee JJ. Second-generation PLINK: rising to the challenge of larger and richer datasets. GigaScience. 2015;4:16.

28. Hamaker EL, Kuiper RM, Grasman R. A Critique of the Cross-Lagged Panel Model. Psychol Methods. 2015;20(1):102–16.

29. Price AL, Patterson NJ, Plenge RM, Weinblatt ME, Shadick NA, Reich D. Principal components analysis corrects for stratification in genome-wide association studies. Nature Genet. 2006;38(8):904–9.

30. Muthen LKM, B. O. Mplus User’s Guide 8th 8ed. Los Angeles, CA: Muthen & Muthen; 1998-2018.

31. Allen HL, Estrada K, Lettre G, Berndt SI, Weedon MN, Rivadeneira F, et al. Hundreds of variants clustered in genomic loci and biological pathways affect human height. Nature. 2010;467(7317):832–8.

32. Hallquist MN, Wiley JF. MplusAutomation: An R Package for Facilitating Large-Scale Latent Variable Analyses in Mplus. Struct Equ Modeling. 2018;25(4):621–38.

33. Team RC. R: A language and environment for statistical computing. (3.3.1). Vienna, Austria: R Foundation for Statistical Computing; 2016.

34. Iacono WG, Malone SM, McGue M. Behavioral disinhibition and the development of early-onset addiction: Common and specific influences. Annual Review of Clinical Psychology. Annual Review of Clinical Psychology. 42008. p. 325–48.

35. Salvatore JE, Savage JE, Barr P, Wolen AR, Aliev F, Vuoksimaa E, et al. Incorporating Functional Genomic Information to Enhance Polygenic Signal and Identify Variants Involved in Gene-by-Environment Interaction for Young Adult Alcohol Problems. Alcoholism (NY). 2018;42(2):413–23.

36. Kichaev G, Bhatia G, Loh PR, Gazal S, Burch K, Freund MK, et al. Leveraging Polygenic Functional Enrichment to Improve GWAS Power. Am J Hum Genet. 2019;104(1):65–75.

37. Lambert SA, Abraham G, Inouye M. Towards clinical utility of polygenic risk scores. Hum Mol Genet. 2019;28(R2):R133–R42.

38. Torkamani A, Wineinger NE, Topol EJ. The personal and clinical utility of polygenic risk scores. Nat Rev Genet. 2018;19(9):581–90.

39. Mostafavi H, Harpak, A., Conley, D., Pritchard, J. K., & Przeworski, M.. Variable prediction accuracy of polygenic scores within an ancestry group.. 2019.

40. Walters RK, Polimanti R, Johnson EC, McClintick JN, Adams MJ, Adkins AE, et al. Transancestral GWAS of alcohol dependence reveals common genetic underpinnings with psychiatric disorders. Nature Neuroscience. 2018;21(12):1656-+.

41. Martin AR, Kanai M, Kamatani Y, Okada Y, Neale BM, Daly MJ. Clinical use of current polygenic risk scores may exacerbate health disparities. Nature Genet. 2019;51(4):584–91.

42. Vrieze SI, Hicks, B. M., Iacono, W. G., McGue, M. Decline in genetic influence on the co-occurrence of alcohol, marijuana, and nicotine dependence symptoms declines from age 14 to 29. American Journal of Psychiatry. 2012;169:1073–81.

43. Dick DM, Rose RJ, Viken RJ, Kaprio J, Koskenvuo M. Exploring gene-environment interactions: Socioregional moderation of alcohol use. Journal of Abnormal Psychology. 2001;110(4):625–32.

